# *In silico* discovery of pathogenic PD-L1 nsSNVs with aberrant glycosylation and immunotherapy binding

**DOI:** 10.1101/2025.06.17.660108

**Authors:** Kai Wei Tan, Shruti Vijay Khare, Joanne Wei En Ong, Carel Min Jie Tan, Sven Hans Petersen, Srinivasaraghavan Kannan, Chandra Shekhar Verma, Ann-Marie Chacko

## Abstract

Immune checkpoint inhibitors (ICIs), particularly anti-PD-L1 monoclonal antibodies, block extracellular interactions between programmed death ligand-1 (PD-L1) and programmed cell death protein-1 (PD-1) to enhance antitumour immunity. Here, we present a streamlined workflow integrating public databases, bioinformatics tools, and *in silico* molecular dynamics (MD) simulations and *in vitro* experimental assays to identify consequential pathogenic nonsynonymous PD-L1 variants (PD-L1^SNVs^). Four variants predicted as pathogenic across six bioinformatic tools (SIFT, PolyPhen-2, PANTHER, SNP-PhD, SNP&GO, and Pmut) destabilized the anti-PD-L1 ICI atezolizumab-drug binding epitope in MD studies, correlating with reduced drug affinity. Live-cell assays directly linked PD-L1^SNVs^ to intracellular trafficking defects and aberrant glycosylation, even distal from *N*-glycosylation sites. Allostery modeling (AlloSigMA) linked these disruptions to long-range structural perturbations. Our work establishes bioinformatics-driven predictions of functionally impactful pathogenic PD-L1^SNVs^ and highlight how glycosylation/trafficking defects may serve as key mechanisms compromising therapeutic efficacy. These insights in predicting and validating pathogenic PD-L1^SNVs^ has significant implications for guiding the personalized selection of effective PD-L1-targeted therapies in clinical settings.

**One Sentence Summary:** Uncovering the implication of non-synonymous single nucleotide variants (nsSNVs) on the structure and function of Programmed Death Ligand-1.

## Introduction

Nonsynonymous single nucleotide variants (nsSNVs) are genetic mutations that alter protein sequences and can potentially affect protein folding, protein-protein interactions (PPI), and even drug-target interactions (*1*). While most nsSNVs are benign, some can lead to a loss or gain of protein function, thereby increasing an individual’s susceptibility to disease. Notably, nsSNVs have been linked to conditions such as hemoglobinopathies (*2*), diabetic nephropathy (*3*) and various cancers (*4, 5*). The association likely stems from the ability of nsSNV to disrupt protein stability, folding, and/or interactions. Furthermore, nsSNVs may allosterically influence distal regions of the protein, potentially resulting in functional changes with broader implications.

Membrane proteins, including the cancer immune check-point proteins programmed death protein 1 (PD-1) and programmed death ligand 1 (PD-L1), constitute a significant portion of cancer drug targets. PD-1 and PD-L1 immune checkpoint inhibitors (ICIs) represent the largest class of immunotherapeutics, which have revolutionised cancer treatment compared to conventional chemotherapy (*6, 7*). However, these treatments show efficacy in only 20% to 40% of patients (*8*). This variable response is largely attributed to differential PD-L1 expression within tumor tissue (*9, 10*). To date, three studies have linked non-protein-altering synonymous SNVs in the 3’ untranslated region (UTR) of the hPD-L1 gene that are associated with treatment outcomes for Nivolumab, an anti-PD-1 monoclonal antibody (mAb) (*11–13*). These mutations influence PD-L1 protein expression within the tumour without altering the protein’s structure. However, the potential role of nsSNVs in PD-L1 (PD-L1^SNVs^) in modulating ICI responses remains unexplored.

In this study, we outline a bioinformatics-driven workflow (Fig 1) utilizing a suite of *in silico* tools to predict pathogenic PD-L1^SNVs^ from general and disease-specific population databases. We further investigated the structural impacts of these mutations relative to the wild-type PD-L1 (PD-L1^WT^). Using atomistic computational modelling/molecular dynamics (MD) simulations, previously applied to understanding PD-L1 protein dynamics (*14, 15*), and *in vitro* assays, we assessed the consequence of PD-L1^SNVs^ on protein structure and their potential impact on the binding of Atezolizumab (Atz), a widely used and clinically approved anti-PD-L1 ICI mAb. By examining how each nsSNV affects critical protein-drug binding interactions, and therefore treatment efficacy, our findings can potentially help identify the most effective drug choices among available options and contribute to precision medicine strategies aimed at designing drugs targeting specific motifs within the PD-L1 protein.

**Fig. 1.**
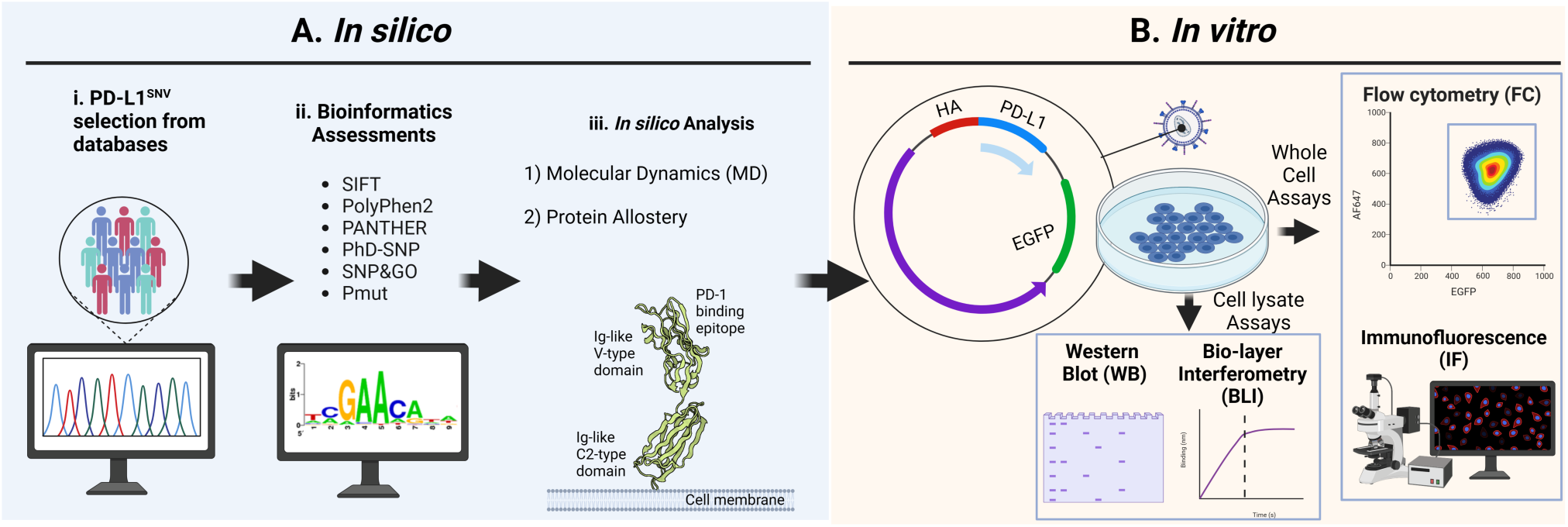
Schematic illustration of the *in silico* and *in vitro* validation workflow for PD-L1^SNV^ missense variants. This approach is divided into two main parts: (**A**) *in silico* and (**B**) *in vitro* validation. (**A**) (**i**) nsSNVs (n = 194) were retrieved from the publicly available UniProt database, and focusing on human variants within the ectodomain and excluding truncating mutations in PD-L1. (ii) Pathogenic nsSNVs were then shortlisted using six bioinformatics tools: SIFT, PolyPhen-2, PANTHER, PhD-SNP, SNP&GO and Pmut. (iii) Shortlisted PD-L1 variants were subjected to *in silico* analyses, including molecular dynamics (MD) simulations and protein allostery predictions. (**B**) Variants were further validated through *in vitro* studies using cellular lysates for, western blot (WB), and biolayer interferometry (BLI), and whole cells for flow cytometry (FC) and immunofluorescence (IF) analyses.

## Results

### Bioinformatics tools identify candidate pathogenic PD-L1^SNVs^

Missense variant data (n=194) was manually curated from Uniprot. To focus on PD-L1^SNVs^ likely to impact anti-PD-L1 ICI binding, we excluded variants within the signal peptide and those causing protein truncation, as they could substantially alter protein structure (Fig. 2A). The remaining nsSNVs (n=180) were analysed using six independently validated, web-based bioinformatic tools that assess the pathogenic potential of protein sequence variants: SIFT (*16*), PolyPhen-2 (*17*), PANTHER (*18*), SNP&GO (*19*), Pmut (*20*) and PhD-SNP (*21*). These tools predict changes in protein physiochemical properties (*16, 17*) tertiary structure (*17, 19*), and residue conservation across protein families based on spatial positions (*16, 17, 19*) and evolutionary history (*18*). SNP&GO, Pmut, and PhD-SNP leverage machine learning algorithms trained on population and clinical datasets to identify pathogenic nsSNVs (*20, 21*).

**Fig. 2.**
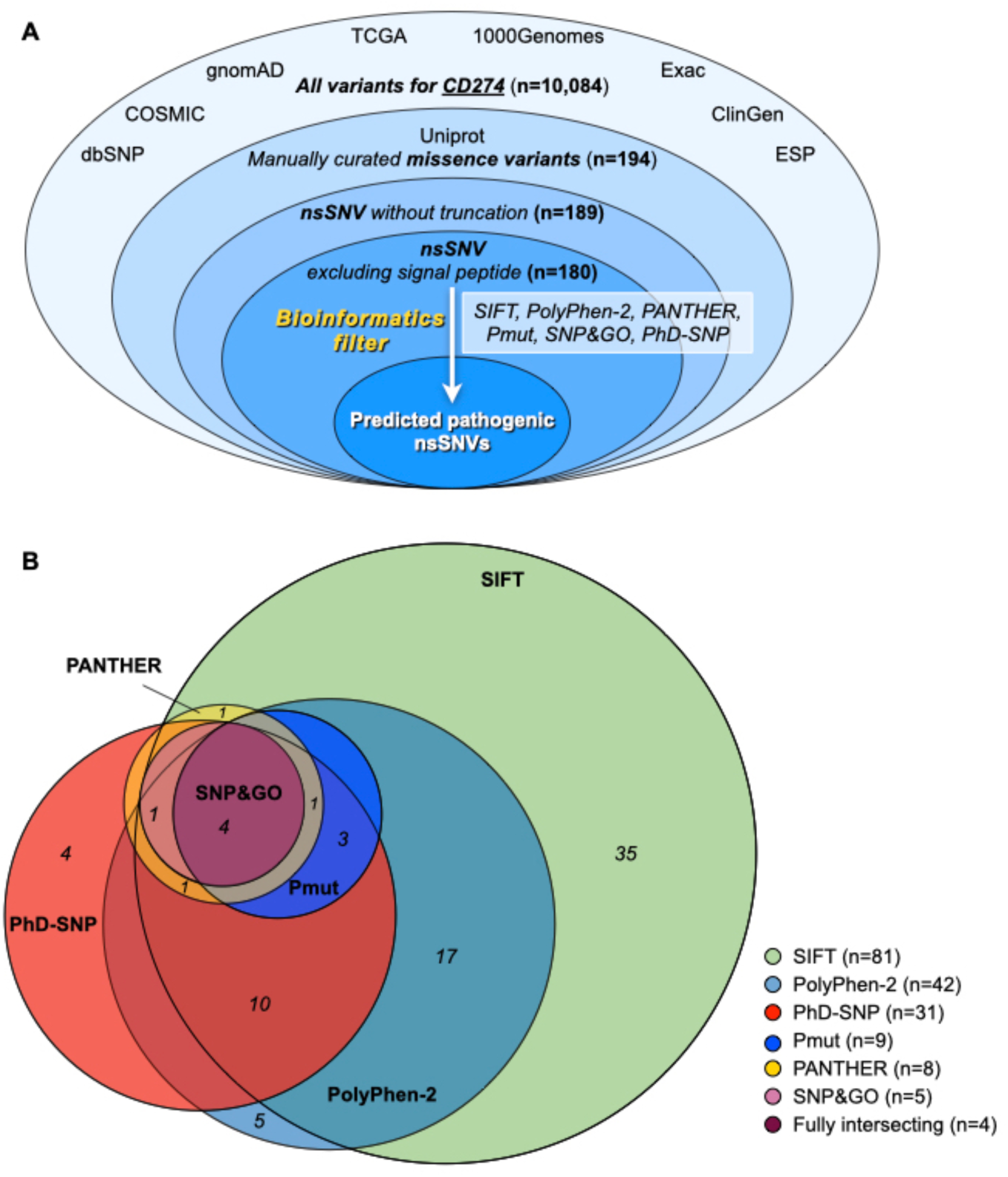
Bioinformatics analysis shortlisted four pathogenic PD-L1^SNVs^ for further experimental validation. (**A**) nsSNVs were obtained from UniProt, and variants located within the signal peptide or resulting in nonsense mutations were excluded. The remaining 180 nsSNVs were subjected to bioinformatics analysis using six tools, with pathogenic variants identified based on specific thresholds:. nsSNVs were shortlisted based on the threshold applied for the respective bioinformatic process. SIFT identified 34 variants as pathogenic (SIFT ≤ 0.05), PolyPhen-2 identified 16 as deleterious (PolyPhen-2 ≥ 1), PANTHER identified 8 variants as pathogenic (PANTHER > 0.5), PhD-SNP identified 31, SNP&GO identified 5, and Pmut identified 9 variants as pathogenic (Pmut > 0.5) (**B**) Four nsSNVs (C40Y, W57C, G159S and W167C) were shortlisted from the intersection of all dix bioinformatics tools (overlapped in the Venn diagrams) using DeepVenn (*45*) for further experimental validation.

Variants were scored for pathogenicity based on the threshold defined by each bioinformatic tool. The SIFT tool identified 45% (n=81) of the 180 curated nsSNVs as deleterious (SIFT ≤ 0.05). PolyPhen-2 flagged 24% (n=42) as probably damaging (PolyPhen-2 > 0.95), while PANTHER identified 8 nsSNVs (4.4%). Pmut, SNP&GO and PhD-SNP shortlisted 9 (5%), 5 (2.8%), and 31 (17.2%) pathogenic nsSNVs, respectively (score = 0.5-1). A complete list of the nsSNVs predicted as pathogenic by the various tools is provided in Table S1. Four variants— C40Y, W57C, G159S and W167C— were identified as pathogenic across all tools, and were hence shortlisted as candidate PD-L1^SNVs^ for validation in a series of experimental tests (Fig. 2B).

### Predicted PD-L1^SNVs^ perturb *in silico* binding to Atz

While the full 3D atomic structure of PD-L1 has not been reported, the crystal structure of its ectodomain containing two Ig-like domains is resolved. The visualization of the hPD-L1 crystal structures (PDB ID: 3BIS, resolution 2.64 Å (*22*)) (Fig. 3) revealed that C40 and W57 are buried within the Ig-like V-type domain of PD-L1. This domain is responsible for binding PD-1 and anti-PD-L1 ICIs such as Atz. Similarly, G159 and W167 are buried within the Ig-like C2-type domain of PD-L1 (Fig. 3A). The accessible surface area (ASA) of the four residues is minimal: 0 Å^2^ for C40 and W57, 0.3 Å^2^ for G159 and 2.2 Å^2^ for W157, which represents less than 1% of the total ASA, confirming that these residues as buried (*23, 24*). Buried residues are crucial for protein folding and structural integrity, so mutations at these positions may destabilize PD-L1’s conformation (*24*). Notably, C40Y, W57C and W167C PD-L1^SNVs^ involve the loss or gain of cysteine residues, potentially leading to aberrant disulphide bridge formation. Specifically, C40Y would disrupt the disulphide bridge between C40 and C114, while W57C could potentially introduce new disulphide bonds between W57C and C40 or C114. Likewise, W167C could potentially disrupt the disulphide bridge between C155 and C209 by forming a new disulphide bond between C167 and C155 or between C167 and C209 (Fig. 3A). The mutation of glycine at position 159 to the polar residue serine could impact PD-L1’s structural flexibility and stability, as glycine is crucial for maintaining protein flexibility (*25*).

**Figure 3:**
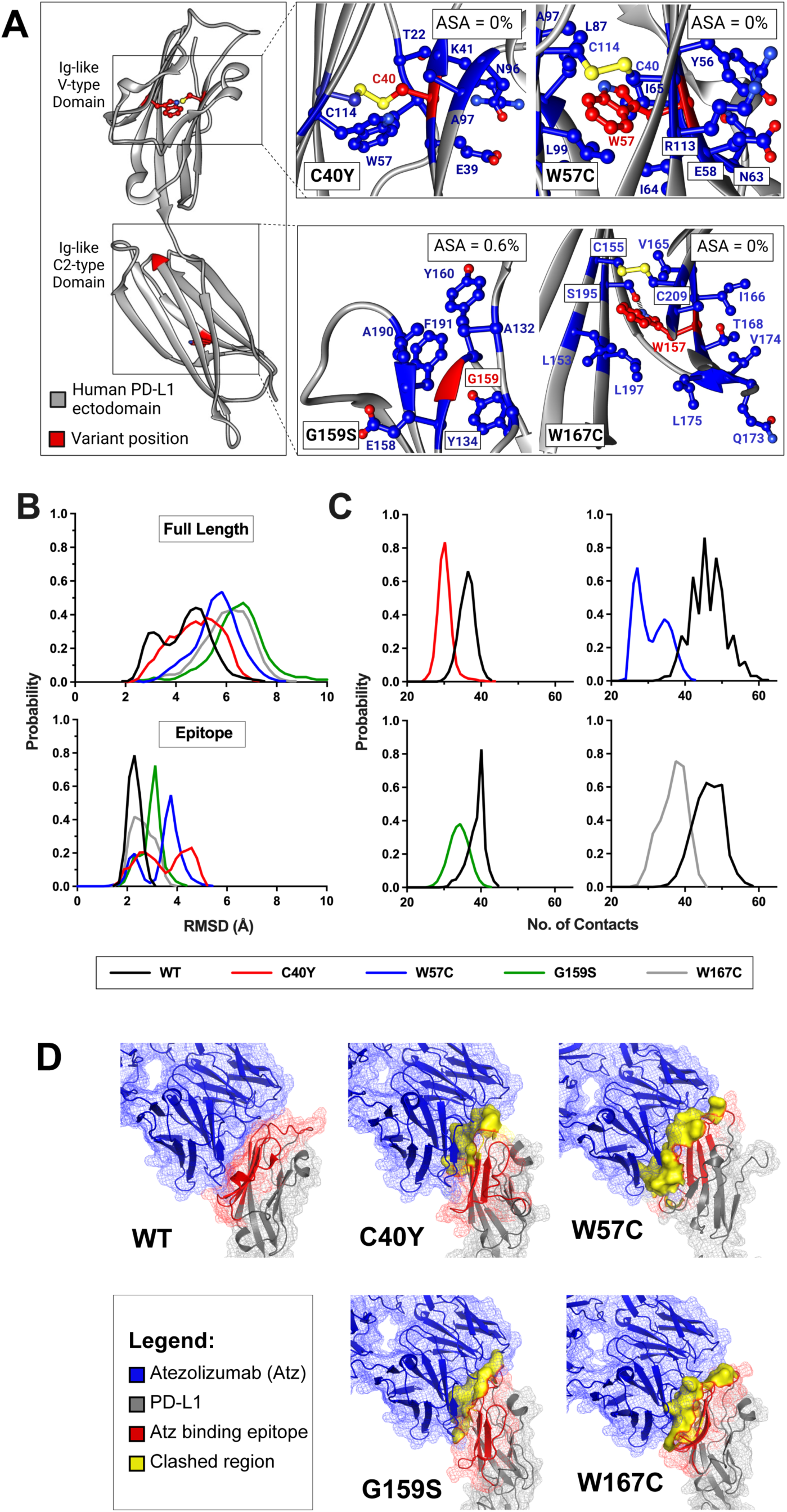
I*n silico* analysis of nsSNVs reveals perturbations in the structural stability of hPD-L1. (**A**) The crystal structure of PD-L1 (green; PDB ID: 3BIS) is shown with the positions of the four nsSNV variants (C40Y, W57C, G159S, W167C) highlighted in red. Residues within a 4Å radium of the variant positions are shown in blue. Accessible surface area (ASA) calculations indicate that all four variant residues are buried (<20% ASA), suggesting that they are located in the interior of the protein structure. (**B**) Distribution of root mean square deviation (RMSD) are shown for the full length PD-L1 (top) and the epitope region (bottom) epitope) sampled during MD simulations of PD-L1^WT^ and PD-L1^SNVs^. (**C**) The number of contacts between each variant residue (C40, W57, G159, W167) and its surrounding residues was quantified during MD simulations. The number of contacts, and the corresponding variant residues are indicated by their respective colors (C40Y:red, W57C: blue, G159S: green,W167C: grey). (**D**) Comparison of the crystal structure of the PD-L1–Atz Fab complex with the conformations sampled during MD simulations of PD-L1^nsSNVs^. Atz Fab is represented as a blue mesh, PD-L1 as a grey mesh, and the Atz binding epitope is shown in red. The conformational changes observed in the in Atz binding epitope of the mutants resulted in steric clash with the antigen-binding site of Atz (yellow surface), disrupting the binding interaction.

We next examined the impact of the mutations on the structure and dynamics of PD-L1. To determine whether PD-L1^SNVs^ cause structural perturbations that could affect binding to Atz, MD simulations were performed on the crystal structure of PD-L1 ectodomains complexed with the Fab domains of Atz (PDB ID: 5XXY, resolution 2.9 Å), which served as the PD-L1^WT^ model. The four mutant structures were generated from this template using Chimera (*26*). The Atz-binding epitope was defined as PD-L1 amino acids 45 to 80 and 112 to 125. Conformational changes in PD-L1 relative to the wild type structure were evaluated used the root mean square deviation (RMSD) of the protein backbone. While the RMSD of the full-length mutants were similar (within 2 Å of each other) relative to PD-L1^WT^ (Fig. 3B, top), the RMSD of the Atz binding epitope varied between 2−5 Å (Fig. 3B, bottom). The most pronounced effect was seen in C40Y, where the substitution of cysteine with the larger tyrosine residue disrupted interactions with neighbouring residues in PD-L1^WT^ (Fig. 3C). Comparison of the PD-L1^SNVs^ conformations (*i.e.,* most populated during simulations) with the experimental PD-L1–Atz Fab complex structure revealed conformational changes in the epitopes of PD-L1^SNVs^, likely resulting in clashes with Atz that prevent effective binding (Fig. 3D).

### *In silico* predicted PD-L1^SNVs^ perturb cell-free PD-L1 binding to Atz *in vitro*

To validate *in silico* binding results, PD-L1^WT^ and PD-L1^SNV^ variants (C40Y, W57C, G159S and W167C) were stably expressed in HEK293T cells using a PD-L1 gene construct with several co-expression tags, including an *N*-terminal haemagglutinin (HA) (fig. S1). As benchmarks, previously reported PD-L1^SNVs^ E58A and Q66A were included. These mutants are considered to be the most disruptive and least disruptive, respectively, for Atz binding (*27*). First, the total PD-L1 expression levels in the PD-L1 isogenic cell lines were assessed by immunoblotting cellular lysates using anti-HA mAb, ensuring that the tag-binding was agnostic of PD-L1 folding status (Fig. 4A and 4B). PD-L1^SNV^ protein expression levels were significantly lower (<50%) compared to PD-L1^WT^ (Fig. 4B, Table S2). Interestingly, there was differential expression of two distinct PD-L1 isoforms — 54 kDa and 47 kDa (Fig. 4A and 4B). The 54 kDa isoform was predominant (>70%) in PD-L1^WT^ and the G159S variant, while the 47 kDa PD-L1 isoform was predominant (>70%) in C40Y, W57C, and W167C variants (Fig. 4B, Table S2).

**Fig. 4.**
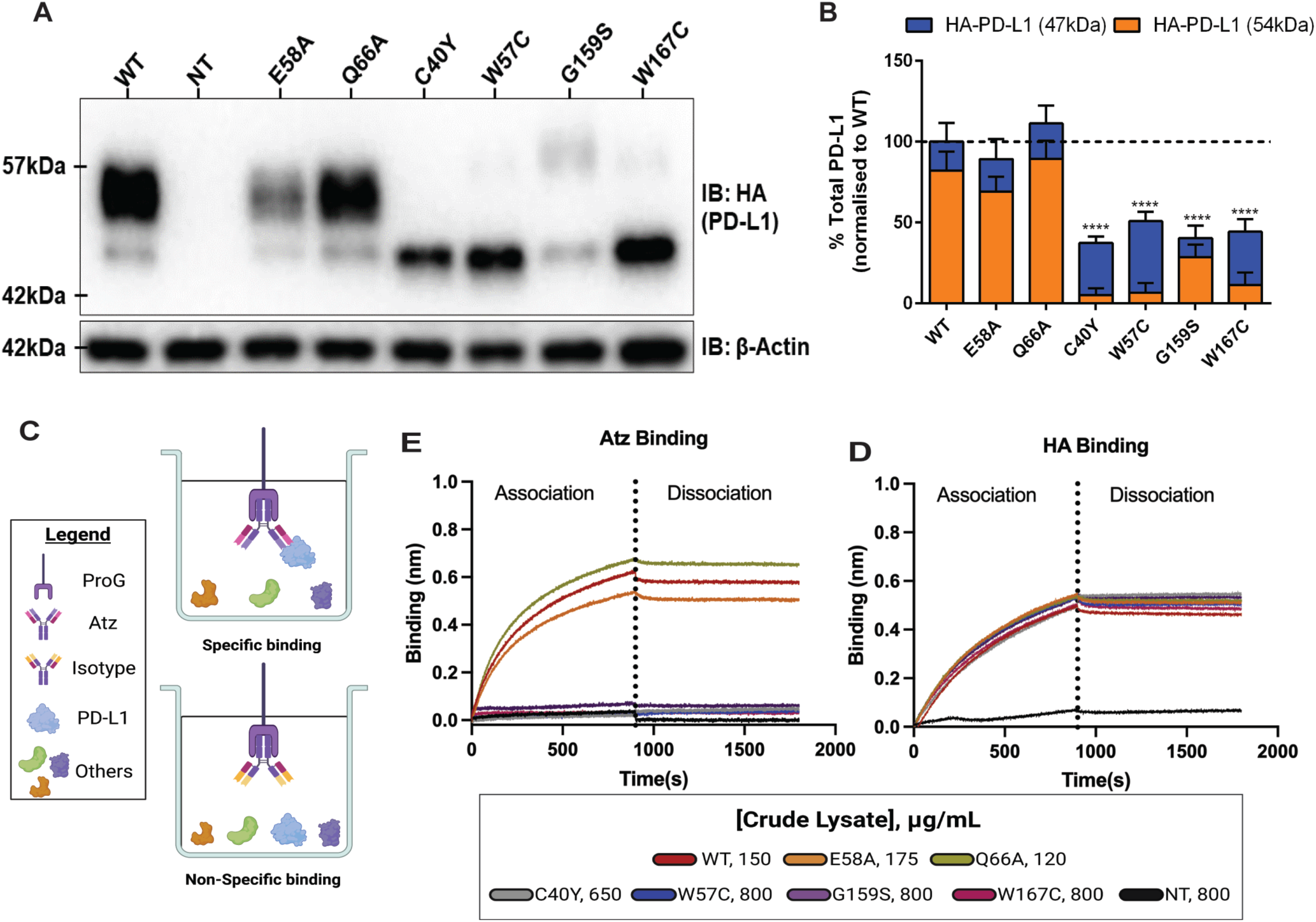
Shortlisted PD-L1^SNVs^ perturb both *in silico* and *cell-free in vitro* binding of anti-PD-L1 ICI therapeutic antibody Atz. (**A**) Immunoblot analysis confirming the protein expression of PD-L1^WT^ and its variants in stably expressing HEK293T cells. (**B**) Quantification of total HA-PD-L1 expression in variants compared to PD-L1^WT^ protein. The fraction of 54 kDa (orange) and 47 kDa (blue) PD-L1 protein isoform bands is shown relative to the total HA-PD-L1. Statistical significance was determined by two-way ANOVA with Sidak’s multiple comparisons test to compare expression of PD-L1^SNVs^ and PD-L1^WT^ (****, *P* < 0.0001). (**C**) Schematic illustrating the set-up for the kinetic binding assay to evaluate the interaction between PD-L1 to Atz using crude lysates from isogenic HEK293T cell lines expressing either PD-L1^WT^ or variants. Specific binding of PD-L1 to Atz was normalised to the non-specific binding of PD-L1 to human IgG_1_ isotype control (right panel). (**D**) HA-PD-L1 protein binding sensogram for crude lysate to ensure equal amounts of HA-protein across lysates. Binding of HA-PD-L1 to anti-HA mAb was normalised using Rabbit IgG isotype control. PD-L1 protein binding sensogram to Atz at crude lysate concentration, with comparable amount of HA-protein. Binding of PD-L1 to Atz was normalised using human IgG_1_ isotype control. Non-transduced (NT) HEK293T lysate served as a negative control to demonstrate the specificity of HA-PD-L1 binding to anti-HA mAb and Atz to PD-L1.

Bio-layer Interferometry (BLI) (*28*) was used to assess the relative binding of Atz to immobilized PD-L1^WT^ or PD-L1^SNVs^ from HEK293T cell-free lysates (Fig. 4C). To ensure consistent loading of PD-L1 variants, crude lysates were titrated before performing Atz binding studies. No binding was detected in lysates from non-transduced (NT) HEK293T cells (Fig. 4D). In experiments with equivalent amounts of HA-tagged PD-L1, PD-L1^WT^, E58A and Q66A bound to Atz (Fig. 4E). Q66A bound to Atz more rapidly than PD-L1^WT^ (k_on_Q66A_ > k_on_WT_), while E58A exhibited slower binding (k_on_E58A_ < k_on_WT_), consisten with previous findings (*27*). Remarkably, none of our four PDL1^SNVs^ variants showed any binding to Atz, with binding profiles resembling those of the NT control.

### PD-L1^SNVs^ compromise membrane trafficking and subvert PD-L1 cellular function

*In silico* experiments predicted that certain PD-L1 mutations could impact its molecular stability, and its binding to Atz in a cell-free context. However, it is unclear how these mutations affect PD-L1 in a more dynamic, live cell environment, particularly with regard to cellular localization. Live-cell flow cytometry (FC) tracking the EGFP reporter co-expressed with PD-L1 confirmed similar transduction efficiencies across all isogenic cell lines (Fig. 5A). Hence, the differential PD-L1 expression levels observed in immunoblotting likely resulted from reduced mutant protein translation or altered trafficking, leading to increased protein degradation due to instability. Live cell FC analysis revealed that surface protein expression of PD-L1^SNVs^ was significantly lower than that of PD-L1^WT^. G159S exhibited the highest surface expression (∼30% of PD-L1^WT^), followed by W57C (∼11%), and W167C (∼4%), while C40Y showed almost no cell surface expression (Fig. 5A, C). Interestingly, the membrane fraction of PD-L1 corresponded with the presence of the 54 kDa PD-L1 band in the immunoblot (Fig. 4A, B), suggesting that only the 54 kDa isoform of PD-L1 is translocated to the membrane. This was further confirmed by permeabilizing cells to detect total PD-L1 (surface and intracellular). Relative to PD-L1^WT^, >90% of HA-tagged PD-L1^SNVs^ was detected in the fixed and permeabilised cells, except for G159S showing 72% of total PD-L1 (Fig. 5A, C). These results suggest that although PD-L1^WT^ and PD-L1^SNVs^ are similarly expressed in total protein levels, the variants are aberrantly trafficked to the cell membrane.

**Fig. 5.**
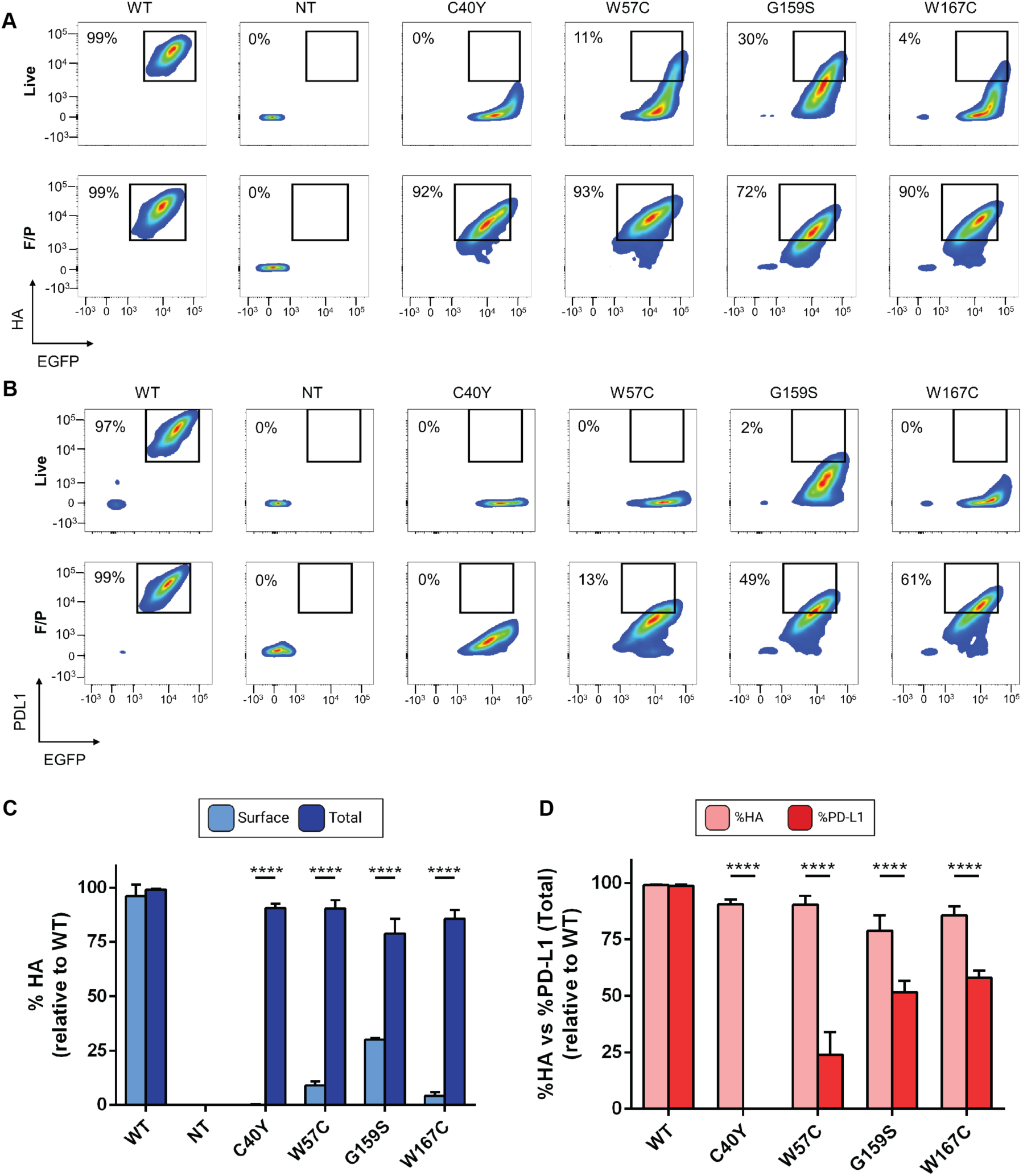
PD-L1^SNVs^ are poorly expressed on the cell membrane. (**A**) Live cell staining for surface PD-L1 expression and fixed/permeabilised (F/P) staining for total PD-L1 expression were performed using anti-HA mAb. Flow cytometry (FC) analysis was conducted to detect surface and total hPD-L1 levels, with the percentage of HA-positive cells (%HA) from PD-L1^SNVs^ normalised with %HA of PD-L1^WT^. EGFP was used as a reporter for hPD-L1 transduction. The FC contour plot highlights the reduced surface PD-L1 expression in the variants compared to PD-L1^WT^. (**B**) Surface and total hPD-L1 expression was further evaluated using anti-PD-L1 mAb by FC. The percentage of PD-L1-positive cells (%PD-L1) in the variants was normalised to the %PD-L1 of PD-L1^WT^. (**C**) The loss of surface PD-L1 expression in the variants was statistically significant and reproducible across three independent experiments. Statistical significance was determined using two-way ANOVA with Sidak’s multiple comparisons test. (**D**) Comparison between %HA and %PD-L1 of total PD-L1 expression across three replicates (n=3) shows that while PD-L1 is still expressed, binding of anti-PD-L1 mAb binding is reduced in the variants. Statistical significance was assessed using two-way ANOVA with Sidak’s multiple comparisons test and one-way ANOVA with Holm-Sidak’s multiple comparisons test for PD-L1^SNVs^ and PD-L1^WT^ expressions. (**, P < 0.01; ***, P < 0.001; ****, P < 0.0001).

To investigate whether the loss of Atz binding in the variants (Fig. 4D, E) resulted from structural alterations, we investigated PD-L1 engagement with an alternative anti-PD-L1 mAb, clone 29E.2A3, which binds an epitope that is distinct from that of Atz. As expected, PD-L1^WT^ bound to mAb 29E.2A3, while all four PD-L1^SNVs^ showed significantly weaker binding in both live and fixed/permeabilised cells (Fig. 5B, D). The results support our hypothesis that these variants result in alterations to the overall structure of PD-L1 and induce allosteric changes that affect the binding of ICIs.

The reduction in variant PD-L1 expression and poor membrane trafficking also point to the possibility of an unfolded protein response (UPR). UPR, resulting from endoplasmic reticulum (ER) stress, could account for the increased protein turnover and the impaired translocation of PD-L1^SNVs^ to the membrane. To identify the subcellular localization of PD-L1^SNVs^, immunofluorescence staining was performed on fixed and permeabilised isogenic HEK293T cells using specific dyes for the ER, golgi apparatus (GA), and plasma membrane (PM). PD-L1 was detected using fluorophore-conjugated anti-HA mAb. As reported previously, PD-L1^WT^ localized predominantly to the PM (Fig. 6A and fig. S4). However, PM localisation was not observed for most of the variants, except for G159S (fig. S2). Anti-HA mAb staining of C40Y, W57C and W167C co-localized strongly with ConA (a marker for ER), showing a Pearson’s Correlation Coefficient (R) > 0.5, indicative of a strong positive correlation (Fig. 6B, C). These variants exhibited stronger ER localization compared to PD-L1^WT^ and G159S, which showed weaker ER localization (R < 0.3). In contrast, G159S localized moderately to the PM (R > 0.3) (Fig. 6B). These data indicate that G159S, which exhibits higher levels of the 54 kDa PD-L1 (predominant in PD-L1^WT^), also shows the highest PM expression (Fig. 4B). Conversely, C40Y, which predominantly expresses the 47 kDa isoform (Fig. 4B), exhibits poor PM trafficking and is mostly retained in the ER (Fig. 6C). Together, these results underscore how PD-L1^SNVs^ can subvert normal cellular trafficking to and function at the plasma membrane, leading to altered localization and impaired drug binding.

**Fig. 6.**
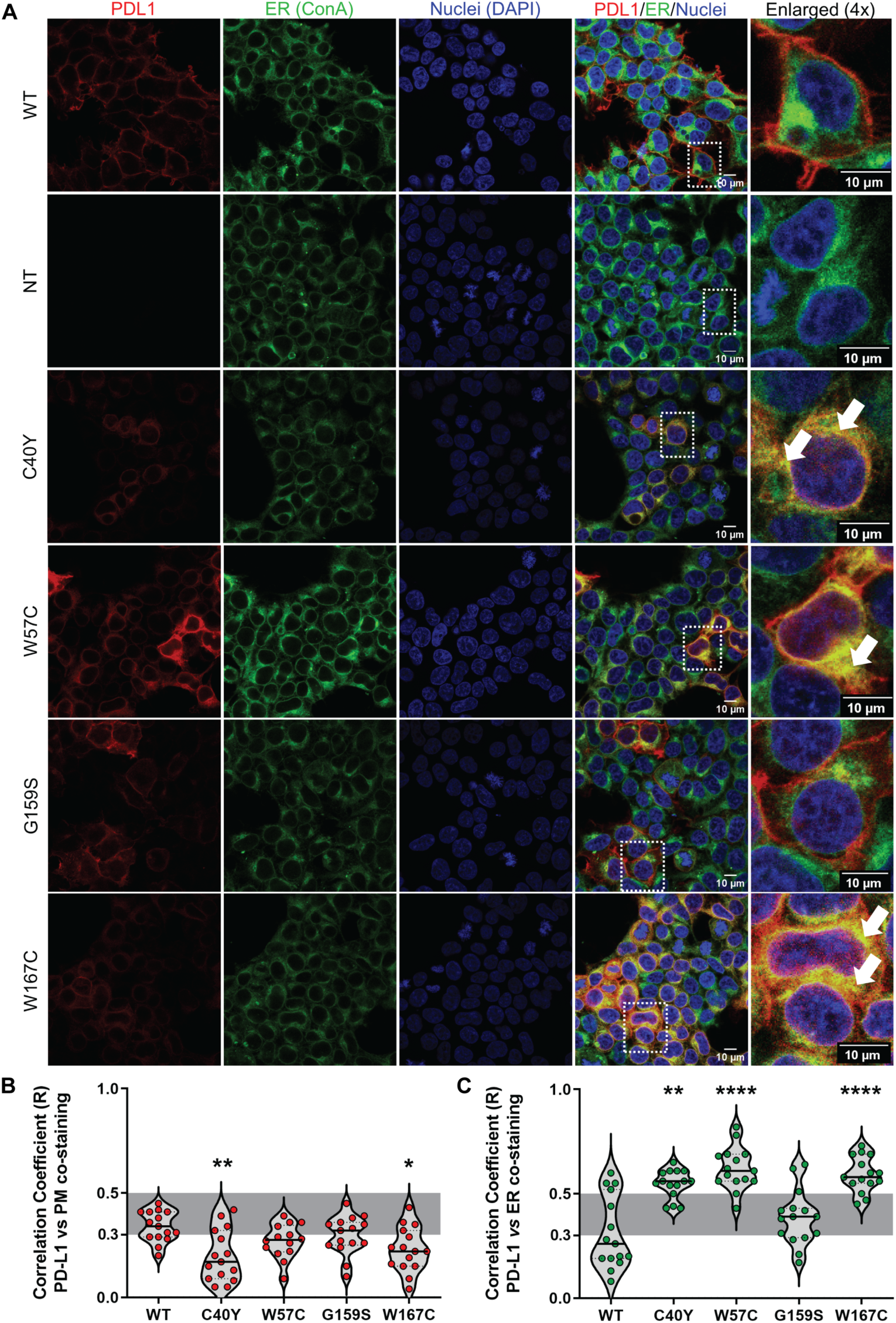
C40Y, W57C and W167C PD-L1^SNV^ variants remain predominantly within the endoplasmic reticulum (ER). (**A**) Immunofluorescence (IF) analysis of PD-L1^WT^ and PD-L1^SNVs^, along with NT control, was performed using the ER marker concanavalin A (ConA; green), anti-HA mAb for hPD-L1 (red) and DAPI for nuclear staining (blue). Colocalization of PD-L1 with the ER marker is indicated by yellow co-staining in the merged/enlarged panels (white arrowhead). (**B, C**) Pearson’s correlation coefficient (*R*) was calculated for the staining patterns of (**B**) hPD-L1 with plasma membrane (PM) marker wheat germ agglutinin (WGA) (as shown in fig. S2) and (**C**) hPD-L1 with the ER marker to assess cellular localization. *R* < 0.3, *R* > 0.3−0.5, and *R* > 0.5 indicated weak, moderate, and strong Pearson’s correlations, respectively. Correlations were determined from three independent experiments, with five selected field of views per experiment (n=15). Statistical significance was calculated using one-way ANOVA with Dunnett’s multiple comparisons test (*, P < 0.05; **, P < 0.01; ****, P < 0.0001).

### PD-L1^SNVs^ disrupt PD-L1 glycosylation

To investigate the source of the size difference between the 54kDa and 47kDa forms of PD-L1, PD-L1^WT^ and PD-L1^SNVs^ protein lysates were de-glycosylated using PNGase F and the results were compared with the glycosylated forms *via* Western blot (fig. S3). As expected, all de-glycosylated PD-L1 proteins were approximately ∼35 kDa, indicating that the primary difference between PD-L1^WT^ and PD-L1^SNVs^ is glycosylation. However, since none of the mutations involve asparagine residues, sites for glycosylation, it is likely that the observed glycosylation differences may be an indirect consequence of structural changes induced by the mutations.

To explore this further, we considered the possibility of allostery, where a mutation can influence the structure and function of regions that are distal from the site of the mutation (*29*). A mutation could alter the size and properties of the side-chain, which in turn might affect packing interactions with neighbouring residues (*30*), potentially propagating to distant sites. To investigate how each nsSNVs may affect glycosylation and binding at the functional site, we used AlloSigMA, a structure-based statistical mechanical model of allostery (SBSMMA). This model first maps the dynamics of the unperturbed protein to understand the allosteric potential of each residue in influencing both neighbouring and distal residues, producing a per-residue allosteric free energy.

The mapping revealed that C40 increases the free energy in residues such as I54, N192 and N200 (blue), while decreasing the free energy in residues D61, N63, R113 and R125 (red) (Fig. 7A). Similarly, the mutations at W57, G159 and W167 impacted both the PD-1/PD-L1 binding epitope and the *N*-glycosylation sites in a similar manner (Fig. 7A). These finding suggests that all four variants could potentially affect both the PD-1/PD-L1 binding epitope and glycosylation sites.

**Fig. 7.**
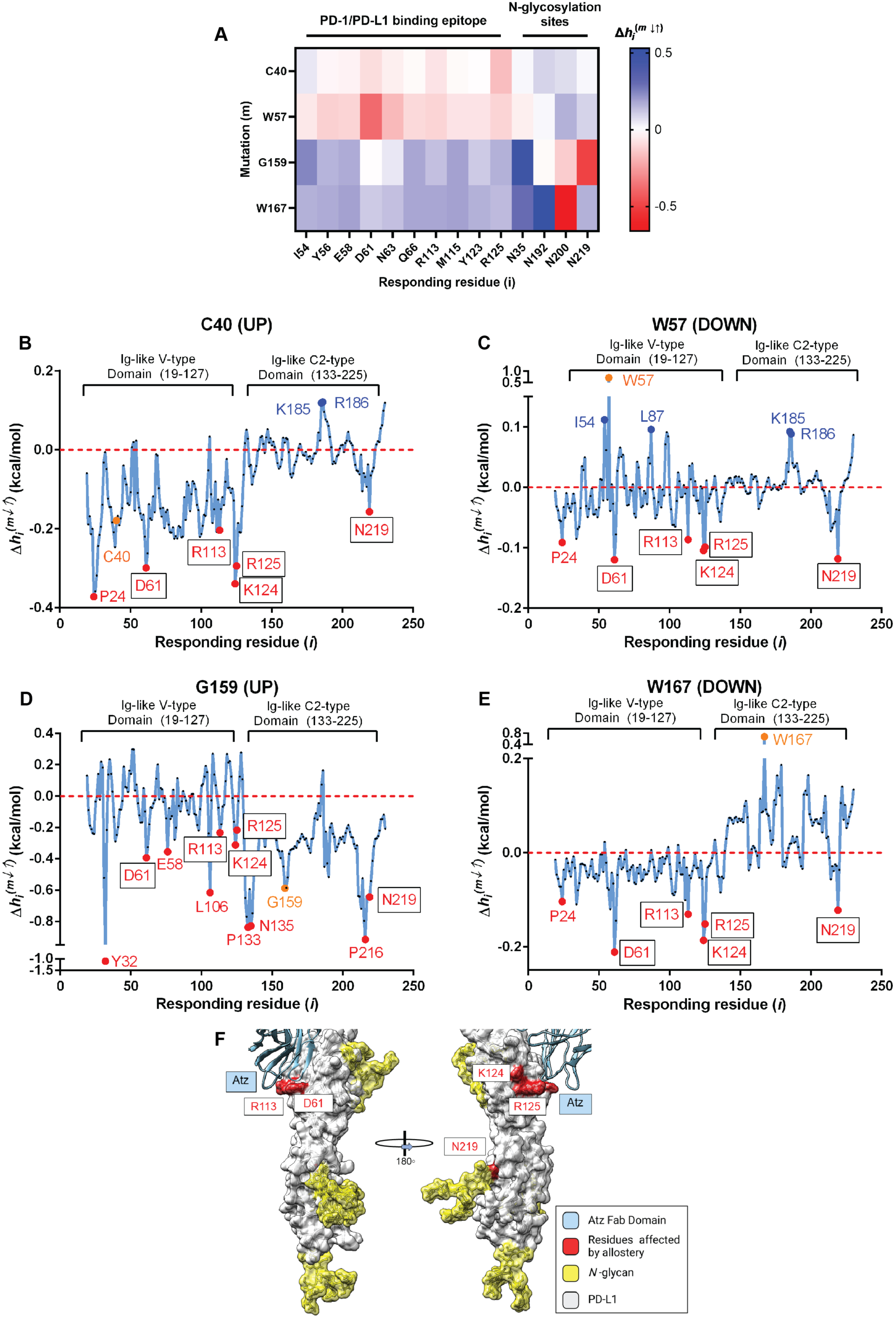
PD-L1^SNVs^ potentially alters the PD-1/PD-L1 binding epitope and *N*-glycosylation sites *via* protein allostery. (**A**) Allosteric Signalling Map (ASM) illustrating the effect of PD-L1^SNVs^ on the PD-1/PD-L1 binding epitope and *N*-glycosylation sites. The characterisation of the amino acid substitution (m) at residues (i) C40, W57, G159 and W167 is presented as Δh_i_^(m↓↑)^, where blue represents a decrease in residue dynamics and red representing an increase. (**B-E**) The residue of interest is indicated in yellow, with residues showing a positive free energy change marked in blue and those residues showing negative free energy change marked in red. Residues consistently affected across all four PD-L1^SNVs^ (D61, R113, K124, R125 and N219) are demarcated with a box. (**B,D**) An UP mutation model was applied to C40 and G159 residues to simulate the substitution with bulkier residues. (**C, E**) A DOWN mutation models was applied to W57 and W167C residues to simulate the substitution with smaller residues. (**F**) Structural representation (merged from PDB ID:5XXY and 3BIK A chain, which includes *N*-glycans) showing how key residues involved in the PD-1/PD-L1 binding epitope (D61, R113, K124 and R125) and the *N*-glycosylation site (N219) are affected by the PD-L1^SNVs^.

To further evaluate the effects of the nsSNVs on the active sites, we employed a separate “Binding Sites and Mutation” module of AlloSigMA. This module simulates the effects of amino acid mutations based on changes in the size of the side chain, referred to as “UP” or “DOWN” mutations, where “UP” mutations estimate an increase in side-chain size (e.g., cysteine to tyrosine), and “DOWN” mutations estimating a decrease side-chain size (e.g., tryptophan to cysteine). For the nsSNVs, UP mutations were assigned to residues C40 and G159 residues, while DOWN mutations were assigned to residues W57 and W167. The results were visualized as per-residue free energy plots (kcal/mol) to assess how each mutation affects the individual residues of the protein.

The analysis revealed that the C40 UP mutation impacted the PD-1/PD-L1 epitope primarily through D61, R113, K124 and R125 and also affected the *N*-glycan site at N219 (Fig. 7B). Similarly, W57 DOWN and W167 DOWN mutations influenced the PD-1/PD-L1 binding epitope via D61, R113, K124 and R125 and also affected the *N*-glycan site at N219 (Fig. 7C and E). In contrast, the G159 UP mutation had a minimal effect on D61, R113, K124, R125 and N219 (Fig. 7D). These results suggest that the C40 UP, W57 DOWN and W167 DOWN nsSNVs predominantly affect the PD-L1 binding epitope through residues D61, R113, K124 and R125, and influence *N*-glycosylation *via* N219 (Fig. 7F). Notably, the G159 UP mutation showed minimal impact of N219 glycosylation, which may explain why the glycosylation pattern of G159S is similar to that of PD-L1^WT^, resulting in the presence of the fully glycosylated ∼54kDa species (fig. S3).

## Discussion

With rapid advances in precision medicine, the customization of treatments based on a patient’s specific genetic background is increasingly becoming a reality. This progress has been driven by the reduction in sequencing costs and an exponential growth in reported genomic variants. However, the comprehensive functional characterization of these variants has not kept pace. While numerous bioinformatics tools, currently around 75 (*31*), are available to identify potentially pathogenic nsSNVs, systematic *in vitro* and/or *in vivo* validation remains scarce. In this study, we present a pipeline that integrates both *in silico* and *in vitro* approaches to predict, test, and validate the impact of PD-L1^SNVs^ on Atz drug binding.

Our study focused on four PD-L1^SNVs^ that were shortlisted using six bioinformatics tools designed to predict structural perturbations from point mutations. While these tools collectively predicted pathogenic PD-L1^SNVs^ with significant structural impact, it remains unclear whether any single tool, or their collective use, can reliably identify pathogenic PD-L1 variants. We hypothesize that the combination of these tools could be generalized for predicting the pathogenicity of variants across various diseases, though it is important to note that the definition of “pathogenicity” can vary depending on the context.

Interestingly, all four of the PD-L1^SNVs^ predicted to perturb PD-L1 protein structure showed significant disruption of Atz drug binding (Fig 4D-E). This was particularly surprising because mutations at G159S and W167C are located on the Ig-like C2-type domain, which is far from the Atz binding epitope situated on the V-type domain (Fig 3A). Our *in silico* studies revealed allosteric effects between the sites of PD-L1 mutations and the Atz binding site, even though the mutations were distal to the epitope (Fig 3B-D). These findings underscore the utility of *in silico* tools, such as MD and AlloSigMA, in elucidating protein dynamics and allosteric interactions. Moving forward, this pipeline could be broadly applied to investigate pathogenic nsSNVs in other protein systems, providing insights into potentially therapeutic targets.

Beyond the allosteric disruptions at the Atz binding epitope, we observed significant alterations in PD-L1 glycosylation patterns and protein expression. Our study revealed that PD-L1 exists as two isoforms: a fully *N*-glycosylated form (54 kDa) and an aberrantly glycosylated form (47 kDa), (Fig 4A, B, Fig. S3). Compared to PD-L1^WT^, the variants C40Y, W57C, and W167C exhibited a marked loss of the fully glycosylated 54 kDa form, with expression levels significantly lower than those of the wild-type (13.1-25.6% vs. 82.2 %, respectively for PD-L1^WT^; Table S2). These findings were unexpected, as the mutations did not involve key asparagine residues (e.g., N35, N192, N200, N219) traditionally required for glycosylation. However, computational analysis suggested that the N219 residue can be significantly affected by these mutations, which could explain the observed differences in glycosylation (Figure 7B, C, E). This result aligns with previous research that has shown that mutations at N219, such as N219Q, result in shorter intracellular half-lives and enhanced degradation of PD-L1, leading to altered glycoform patterns (*32*).

The G159S variant, by contrast, displayed a glycosylation pattern that was similar to that of PD-L1^WT^, though with lower overall protein expression levels (Fig 4B, S3, Table S2). Our *in silico* allostery analysis also suggested that the N219 is less affected in G159S compared to the other variants (Fig 7D), which could explain why G159S maintains a glycosylation profile closer to PD-L1^WT^. This suggests that an alternative degradation pathway, distinct from the one responsible for the degradation of the aberrantly glycosylated variants C40Y, W57C, and W167C, may be at play for G159S. With respect to subcellular localization, flow cytometry (FC) and immunofluorescence (IF) studies confirmed that the C40Y, W57C, and W167C variants predominantly localized to the endoplasmic reticulum (ER), with minimal expression at the plasma membrane (Fig. 5, 6). In contrast, G159S showed a pattern of localization more akin to PD-L1^WT^. A previous study also observed that the aberrantly glycosylated S195E PD-L1^SNV^ trafficked primarily to the ER (*33*). This reduction in fully glycosylated PD-L1 expression at the tumor cell surface, which can also be induced by metformin (33), enhances CD8^+^ T cell infiltration and reduced tumor growth. Taken together, these findings underscore the essential role of *N*-glycosylation in regulating PD-L1 stability and trafficking, which directly influences immune checkpoint modulation. A schematic depicting our proposed models for the cellular fate of PD-L1^nsSNVs^ is shown in Figure 8.

**Figure 8.**
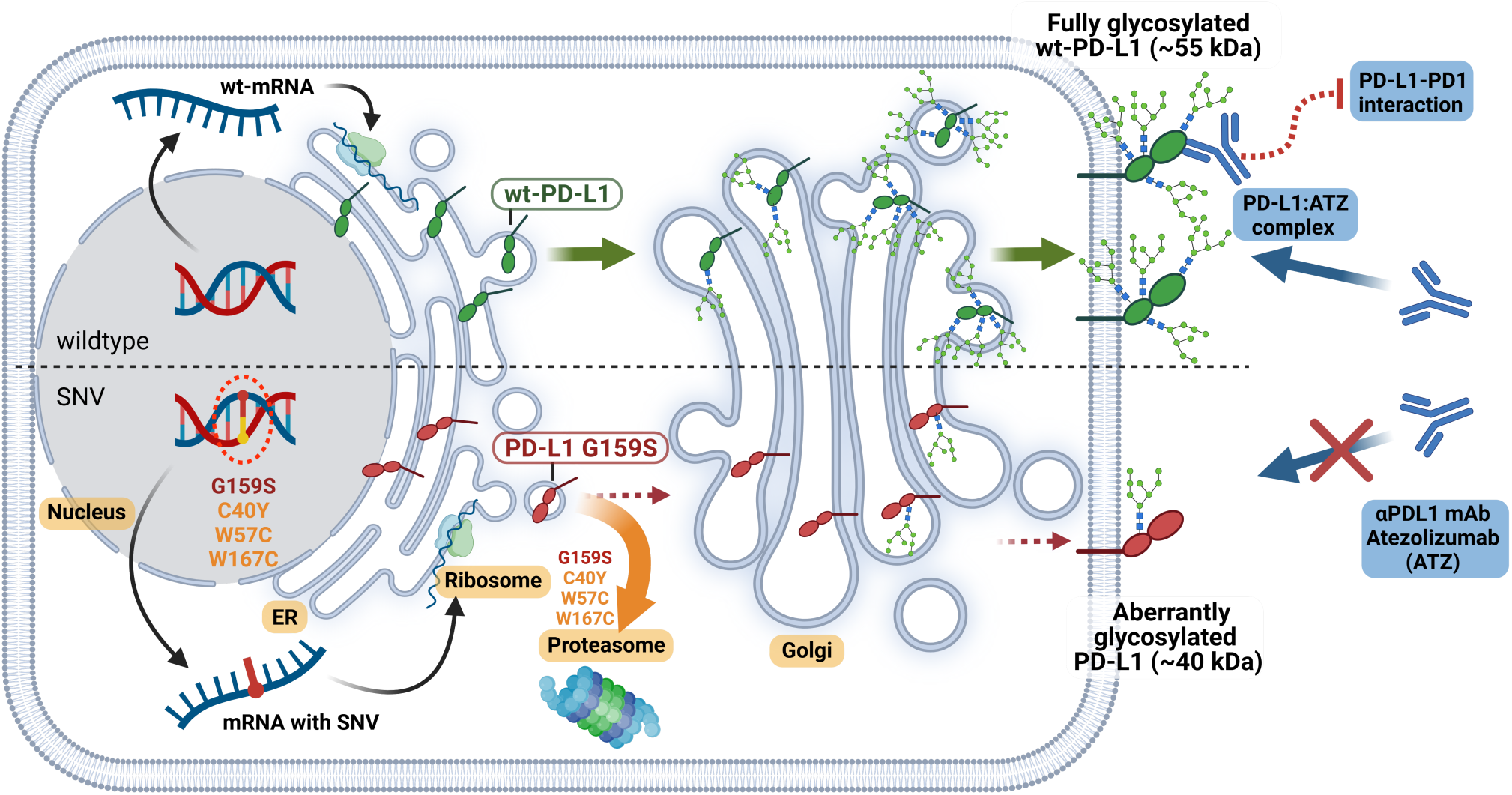
Model of cellular fate of PD-L1 single nucleotide variant (PD-L1^SNVs^) due to aberrant protein glycosylation, membrane translocation, and overall protein turnover. C40Y, W57C, and W167C resulted in a significant loss of the fully glycosylated 54 kDa PD-L1 isoform, which corresponds to a reduction in membrane translocation. This aberrant glycosylation likely causes protein misfolding and retention in the endoplasmic reticulum (ER). This misfolding is associated with increased protein turnover, likely due to ER stress, and enhanced proteasomal degradation. The G159S variant causes a partial loss of the fully glycosylated 54 kDa species, but some portion of the protein still manages to translocate to the cell membrane. The G159S variant results in less severe glycosylation defects compared to the other SNVs, leading to partial cell membrane localization. Overall, aberrant glycosylation caused by these SNVs leads to a loss of functional PD-L1 on the cell surface, increased protein degradation, and impaired membrane trafficking. These effects can alter the protein’s interaction with immune checkpoint inhibitors, potentially influencing therapeutic outcomes.

One limitation of our study is the potential destabilisation of PD-L1^SNVs^ upstream of post-translational modifications, such as *N*-glycosylation. Since G159S exhibited a glycoform expression pattern similar to that of PD-L1^WT^ but at lower protein levels, it suggests the possibility of a competing degradation pathway between glycosylated and aberrantly glycosylated PD-L1 isoforms. Further studies are warranted to investigate the kinetics of these degradation pathways and how they influence the stability and trafficking of PD-L1 variants.

In conclusion, our study provides a comprehensive workflow that integrates *in silico* and *in vitro* approaches to investigate the functional impact of nsSNVs on protein structure. We identified four PD-L1 variants that significantly perturb protein structure, impacting both Atz drug binding and *N*-glycosylation, with downstream effects on protein expression and trafficking. Our approach offers a framework for investigating the consequences of nsSNVs in other protein systems, potentially shedding light on their role is disease pathogenesis and therapeutic resistance.

## Materials And Methods

### Study design

We developed a bioinformatics-driven workflow to predict and test pathogenic PD-L1^SNVs^ *in silico* and validate their molecular impact on both protein structure and function *in vitro*. First, we compiled all reported missense variants in PD-L1 from the curated UniProt database. Using a combination of six bioinformatics tools, we identified four pathogenic nsSNVs. These tools selected deleterious variants in highly conserved residues (*via* SIFT (*16*), PolyPhen-2 (*17*), SNP&GO (*19*) and PANTHER (*18*)) and incorporated features recognized to be pathogenic in machine learning-based algorithms (PhD-SNP (*21*) and Pmut (*20*)). These variants were further analysed using *in silico* methods, such as Molecular Dynamics (MD) simulations to identify structural changes (*34*) and AlloSigMA (*35*) to assess potential functional consequences. Experimental validation included the assessment of protein expression, post-translational modifications, trafficking, and binding of these variants.

### Prediction of pathogenic PD-L1 variants

A dataset of amino acid substitutions resulting in nsSNVs in human PD-L1 (hPD-L1) was obtained from Uniprot using the search term *CD247,* the gene encoding PD-L1 protein. UniProt includes variant data from various databases, including dbSNP (*36*), COSMIC (*37*), gnomAD (*38*), and TCGA. From an initial list of 194 missense variants, we excluded those within the signal peptide (1^st^ to 17^th^ residues) (n=9) and those resulting in a premature stop codon (n=5) (Fig. 2). The remaining PD-L1^SNVs^ (n=180) were further filtered using six independent bioinformatic methods, with pathogenic nsSNVs identified on tool-specific scores: SIFT ≤ 0.5, PolyPhen-2 ≥ 0.95, PANTHER ≥ 0.5, SNP&GO ≥ 0.5, Pmut ≥ 0.5 (*20*) and PhD-SNP ≥ 0.5 (Table S1).

### Structure preparation

X-ray crystal structures of human PD-L1^WT^ protein were downloaded from the Protein Data Bank (PDB): the PD-L1 ectodomain dimer (PDB ID:3BIS, resolution 2.64 Å (*22*)); and the PD-L1 Ig-like V-type domain/Atz Fab domain complex (PDB ID: 5XXY, resolution 2.9 Å (*27*)). These structures were visualized using UCSF Chimera software (v1.10). Point mutations corresponsing to the nsSNVs of interest were introduced into the PD-L1/Atz complex structure using the Dunbrack 2010 rotamer library (*39*) available in Chimera.

### Molecular dynamics simulations

To investigate the conformational changes resulting from the introduction of the PD-L1^SNVs^, molecular dynamics (MD) simulations were carried out using the AMBER programme (*40*). Both wild type and mutant PD-L1 proteins (generated in Chimera using PDB ID: 5XXY) were prepared for MD simulations according to standard protocols (*40*). The systems were represented using the Amber14SB force field (*41*), and sodium and chlorine counterions were added to neutralize the charge of each system simulated. A water box (TIP3) in a truncated octahedron geometry was placed around each protein, with a 10 Å minimum away from the protein. Systems underwent 1000 cycles of minimization through, with 500 steps of steepest descent followed by conjugate-gradient algorithms. The MD simulations were performed with a 2 fs time step at 300 K and 1 atm pressure in the NPT ensemble, applying periodic boundary conditions for an initial equilibration of 2 ns, followed by a 500 ns production run. During each MD run, the average potential energy (EPTOT) and average dihedral term potential energy (DHE) of PDL1^WT^ and its variants were recorded to generate the parameters (i.e., EthreshP, AlphaP, EthreshD, AlphaD) for accelerated Molecular Dynamics (aMD) (*42*). Three independent 500 ns aMD runs were performed for PDL1^WT^ and PDL1^SNVs^. The Particle Mesh Ewald method (*34*) was used electrostatic calculations with a 9 Å cut-off for non-bonded interactions. Isotropic position scaling and a Langevin type thermostat were used to maintain the temperature at 300 K. RMSD and Number of Contacts calculations were performed using the PTRAJ module in Amber18, with contacts defined residues within 6.5 Å of the residue of interest. The VMD (Visual molecular dynamics) program (*25*) and Pymol (*43*) were used to visualize and prepare figures from the MD trajectory.

### Protein allostery

Protein allostery predictions were performed *in silico* using the AlloSigMA webserver (*35*). To test the impact of variants on protein allostery, the hPD-L1 structure (PDB ID: 3BIS (*22*)) was analyzed using the ‘Protein Interfaces, Surfaces, and Assemblies’ (PISA) module to compute the per-residue effects of the variant residues on the protein structure. Allosteric signalling maps were generated using UP and DOWN modulation models (*m↓↑*), reflects the effect of a side-chain size being increased (e.g., cysteine to tyrosine) or decreased (e.g., tryptophan to cysteine), respectively. Although AlloSigMA cannot directly mutate a residue to the variant amino acid, it simulates the effects of UP (bulkier) and DOWN (smaller) mutations. Consequently, UP mutations were applied for the C40Y and G159S variants, and DOWN mutations for the W57C and W167C variants. The change in free energy (Δh_I_) for individual residue mutations was calculated based on the equation below, with the modulation ranging from 1 to −1 kcal/mol:

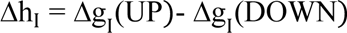

To further assess the allosteric effects, reported Atz binding site residues (I54, Y56, E58, D61, N63, Q66, R113, M115, Y123, K124 and R125) and *N*-glycan sites (N35, N192, N200 and N219) were included (*27,33*). Based on these sites and the mutations selected, a graphical representation was generated to visualize how the UP or DOWN mutations at the residue of interest affected the Atz binding epitope and *N*-glycan sites of PD-L1.

### Cell culture, antibodies, and reagents

The following antibodies were used in this study: for BLI anti-PD-L1 IgG_1_ mAb Atezolizumab (Atz; Tencentriq), human IgG_1_ isotype control (Bio X Cell, BP0297), rabbit IgG anti-HA mAb (CST, C29F4) and rabbit IgG isotype control (Invitrogen, 26102); anti-PD-L1 polyclonal antibody (pAb) (Invitrogen, PA5-28115), and mouse anti-β-actin mAb (Proteintech, 66009-1-ig) as primary antibodies; Secondary antibodies for immunoblotting included goat anti-rabbit horseradish peroxidase (HRP)-conjugated pAb (Proteintech, SA00001-2), and goat anti-mouse HRP-conjugated affinipure pAb (Proteintech, SA000001-1). For flow cytometry (FC) and immunofluorescence (IF), mouse IgG_1_ and anti-HA AF647 conjugated mAb (CST, 6E2) were used. The PhenoVue Cell Painting Kit (Perkin Elmer, PING 11), containing PhenoVue Fluor 488-concanavalin A (ConA), PhenoVue Fluor 555-wheat germ agglutinin (WGA) and PhenoVue Dye diluent A, were used for subcellular staining and fluorescence imaging.

### Generation of isogenic cell line pairs expressing PD-L1^WT^ and PD-L1^SNV^ variants

The PD-L1^WT^ sequence (Gene ID: 29126) and PD-L1^SNV^ variants (C40Y, W57C, G159, W167C, E58A, and Q66A) were cloned into a proprietary lentiviral vector (pD2109-Efs: EF1a-ORF, Lenti-ElecD, ATUM). Each PD-L1 construct included the signal peptide (residues 1-18), extracellular domain (residues 19-290), transmembrane domain (residues 240-261), and intracellular domain (residues 261-291), with a HA tag downstream of the *N*-terminal signal peptide and a GSGSGS linker peptide followed by a BirA biotinylation Avi-tag at the *C*-terminus (fig. S1). EGFP was included as a fluorescent reporter gene. Lentiviral particles were prepared by co-transfecting HEK293T cells (ATCC, CRL-3216) with third generation packaging (pRSV-REV, pMDLg/pRRE; Addgene), envelope (pMD2.G; Addgene) and expression vectors using the transfection enhancer Fugene HD (Promega, TM328) according to manufacturer’s instructions. Virus was harvested 48 h and 72 h post-transfection. Purified viral supernatant was used to transduce fresh HEK293T cells in the presence of polybrene (8 μg/mL; Sigma-Aldrich, 107689). Transduced cells were selected using puromycin (5 µg/mL; Invivogen, ant-pr-1), yielding a transduction efficiency of 90-95 %. Cells were grown in Dulbecco’s Modified Eagle’s Medium (DMEM; Fisher Scientific, 2-800-017) supplemented with 10 % foetal bovine serum (FBS; Hyclone, SV30160.03) and maintained at 37 °C and 5 % CO_2_ in a humidified incubator and used for subsequent studies. Cells were subsequently characterised by immunoblotting, flow cytometry (FC), and immunofluorescence (IF) (*supplemental methods*).

### Kinetic binding assays to PD-L1

Kinetic binding assays for Atz binding to PD-L1 and its variants were perfromed using the Octet Red 96e bio-layer interferometry (BLI) system (Sartorius ForteBio) with 8-channel detection mode and fresh Protein G (ProG) sensors (Sartorius, 18-5082). Assays were conducted in pre-filtered PBST (0.22 µm-filtered PBS (Axil Scientific, BUF-2040) supplemented with 0.05% Tween (Bio-Rad, 1706531)) at 30 °C with orbital shaking at 1,000 rpm. The optimal concentrations of crude lysate corresponding to an equivalent amount of HA-tagged PD-L1 protein was first determined. For each HA binding assay step, the biosensor was sequentially immersed the following solutions (200 µL/well): i) PBST, hydration (10 min), ii) PBST, baseline (60 s), iii) 33 nM anti-HA (CST, C29F4), loading (30 s), iv) PBST, washing (60 s), v) crude lysates (150 µg/mL WT, 175 µg/mL E58A, 120 µg/mL Q66A, 650 µg/mL C40Y, and 800 µg/mL W57C, G159S and W167C), association (15 min), and vi), PBST, dissociation (15 min). Rabbit IgG isotype control (33 nM) loaded on the ProG sensor during step iii) and processed identically was used as the reference. After determining the optimal lysate concentrations, Atz binding to each crude lysate at these concentrations was evaluated. The following steps were performed for the Atz binding experiment: i) PBST, hydration (10 min), ii) PBST, baseline (60 s), iii) 33 nM Atz loading (30 s), iv) PBST, washing (60 s), v) crude lysates at previously determined concentrations, association (15 min), and vi) PBST, dissociation (15 min). Human isotype control IgG_1_ (33 nM) was used as the reference during the loading step (iii). Binding sensorgrams were acquired and processed using ForteBio acquisition (v.10.0.1.3) and analysis (v.10.0) software as previously described (*44*). The HA and Atz binding values were first normalized by subtracting the respective reference values, then aligned to association.

### Statistical analysis

Statistical analyses were performed using GraphPad Prism 6 software. A Mann Whitney U-test was applied for all the experiments unless otherwise noted, with a *P* value threshold of < 0.05 considered statistically significant.

## Acknowledgments

Thanks to Professor Daniel S.W. Tan at the National Cancer Centre Singapore (NCCS) for providing Atezolizumab. Special thanks to Dr. Igor Berezovsky (BII A*STAR) for helpful discussions on AlloSigMA. The PhenoVue^TM^ cell painting kit for immunofluorescence studies was a generously provided by Revvity (Singapore) and Genomax (Singapore). W57C, G19S and W167C PD-L1^SNVs^ were identified from the TCGA Research Network (https://www.cancer.gov/tcga). We also extend our appreciation to Ms. Ravinuthula Sruthi Jagannathan (Duke-NUS) for her editorial assistance.

## Author contributions

Conceptualization: KWT, AMC

Methodology: KWT, SVK, AMC

Analysis: KWT, SVK

Curation of *in silico* data: SVK, SK, CV

Stable cell line generation: SHP

*In vitro* experiments and analysis: KWT, JWEO, CMJT

Resource provision: AMC, CV

Supervision: KWT, AMC

Writing – original draft: KWT

Writing – review & editing: KWT, AMC, SVK, SHP, CMJT, SK, CV

## Data and materials availability

All data are available in the main text or the supplementary materials.

